# Synaptic vesicle release regulates pre-myelinating oligodendrocyte-axon interactions in a neuron subtype-specific manner

**DOI:** 10.1101/2024.01.25.577297

**Authors:** James R. Gronseth, Heather N. Nelson, Taylor L. Johnson, Taryn A. Mallon, Madeline R. Martell, Katrina A. Pfaffenbach, Bailey B. Duxbury, John T. Henke, Anthony J. Treichel, Jacob H. Hines

## Abstract

Oligodendrocyte-lineage cells are central nervous system (CNS) glia that perform multiple functions including the selective myelination of some but not all axons. During myelination, synaptic vesicle release from axons promotes sheath stabilization and growth on a subset of neuron subtypes. In comparison, it is unknown if pre-myelinating oligodendrocyte process extensions selectively interact with specific neural circuits or axon subtypes, and whether the formation and stabilization of these neuron-glia interactions involves synaptic vesicle release. In this study, we used fluorescent reporters in the larval zebrafish model to track pre-myelinating oligodendrocyte process extensions interacting with spinal axons utilizing in vivo imaging. Monitoring motile oligodendrocyte processes and their interactions with individually labeled axons revealed that synaptic vesicle release regulates the behavior of subsets of process extensions. Specifically, blocking synaptic vesicle release decreased the longevity of oligodendrocyte process extensions interacting with reticulospinal axons. Furthermore, blocking synaptic vesicle release increased the frequency that new interactions formed and retracted. In contrast, tracking the movements of all process extensions of singly-labeled oligodendrocytes revealed that synaptic vesicle release does not regulate overall process motility or exploratory behavior. Blocking synaptic vesicle release influenced the density of oligodendrocyte process extensions interacting with reticulospinal and serotonergic axons, but not commissural interneuron or dopaminergic axons. Taken together, these data indicate that alterations to synaptic vesicle release cause changes to oligodendrocyte-axon interactions that are neuron subtype specific.

## Introduction

Oligodendrocytes are central nervous system (CNS) glia that myelinate axons and perform numerous myelin-independent functions (Xiao and Czopka, 2023). Approximately 5% of CNS cells are oligodendrocyte progenitor cells (OPCs), which are motile, multi-polar cells found diffusely throughout the brain and spinal cord (Dawson et al., 2003). In addition to their capacity to differentiate into myelinating oligodendrocytes, OPCs participate in axon guidance, circuit development and refinement, and in some instances, form nonsynaptic junctions and synaptic contacts with neurons (Bergles et al., 2000; Wake et al., 2015; Xiao et al., 2022). Upon differentiation, OPCs undergo a dynamic series of changes to cell morphology and behavior (Baumann and Pham-Dinh, 2001). OPC maturation into premyelinating oligodendrocytes involves the ramification of process extensions into a larger arbor of branchy and exploratory processes (Hardy and Friedrich, 1996). Pre-myelinating oligodendrocyte process extensions presumably contact and sample numerous putative target axons prior to forming myelin sheaths (Almeida and Macklin, 2023; Hill and Grutzendler, 2019; Schnädelbach et al., 2001).

Oligodendrocyte process extensions have the capacity to distinguish between the diverse CNS cell types and their subcellular compartments. Oligodendrocytes selectively enwrap and myelinate neuronal axons despite their morphological similarity to dendrites and glial process extensions (Lubetzki et al., 1993). The molecular pathways used by oligodendrocyte processes to distinguish axonal and somatodendritic domains are beginning to be elucidated (Díez-Revuelta et al., 2017; Klingseisen et al., 2019; Lee et al., 2012; Redmond et al., 2016). Furthermore, oligodendrocytes selectively myelinate specific neural circuits and axon subtypes during development, adaptive myelination, and remyelination (Bacmeister et al., 2022, 2020; Koudelka et al., 2016; Neely et al., 2022; Orthmann-Murphy et al., 2020; Tomassy et al., 2014; Yang et al., 2020). Collectively, these studies point toward a model whereby oligodendrocytes can detect the biophysical profile and molecular cues presented by objects they encounter to determine when and where to form myelin sheaths.

Synaptic vesicles are traditionally viewed as a specialized subset of vesicles restricted to distal axon synaptic terminals. However, synaptic vesicles are also present within proximal axon segments. Axonal synaptic vesicles may be motile or static, and both evoked and non-evoked release can occur along the length of the axons (Kukley et al., 2007; Ziskin et al., 2007). One function of synaptic vesicles in proximal axon segments involves the control of myelin sheath growth. Synaptic vesicles are enriched at sites of active myelination, and positively regulate myelin sheath stabilization and growth along the length of axons (Almeida et al., 2021; Hines et al., 2015; Hughes and Appel, 2019; Koudelka et al., 2016).

Synaptic vesicle release along developing axons occurs most frequently at myelin sheath-axon contact sites, but also occurs at non-myelinated axon segments (Almeida et al., 2021). Whereas synaptic vesicle release stabilizes myelin sheath-axon interactions, how synaptic vesicle release regulates oligodendrocyte process-axon interactions prior to initial axon wrapping and myelination is unknown. In this study, we tested the hypothesis that synaptic vesicle release influences the formation and stabilization of oligodendrocyte-axon interactions prior to myelin sheath formation. Using the larval zebrafish spinal cord model, we found that synaptic vesicle release is involved in establishing the density that pre-myelinating oligodendrocyte processes interact with axons of different neuronal subtypes. Synaptic vesicle release blockade also regulated the temporal dynamics of pre-myelinating oligodendrocyte process interactions with *phox2b*^+^ reticulospinal axons, but this was only observed for process extensions interacting at axon varicosities. In accordance with this observation, synaptic vesicles were enriched at axon varicosities in comparison to intervening axon segments. Together, these data indicate that synaptic vesicle release influences both the spatial and temporal profile of oligodendrocyte-axon interactions.

## Results

To assess the role of synaptic vesicle release in oligodendrocyte-axon interactions, we first performed time-lapse imaging to track dynamic cell-cell interactions in the larval zebrafish spinal cord. We used the Gal4-UAS system to scatter-label individual myelin-competent *phox2b*^+^ reticulospinal axons that projected longitudinally across in the ventral spinal cord with membrane-tethered EGFP. Combined use of the *Tg(sox10:mRFP)* reporter line to label oligodendrocyte process extensions revealed sites of fluorescence colocalization between oligodendrocyte process extensions and reticulospinal axons (Fig. 1A). As a means to preferentially observe pre-myelinating oligodendrocyte process extensions, we performed time-lapse imaging at a developmental stage when *sox10*:mRFP^+^ oligodendrocytes possessed branchy and ramified process extensions, with some forming their first nascent myelin sheaths at the time and position of viewing. During the 90 minute time-lapse imaging period, the majority of oligodendrocyte process extensions had a thin and branchy morphology and had yet to initiate axon wrapping or myelination (Fig. 1A). A subset of *sox10*:mRFP^+^ processes showed interaction with EGFP^+^ reticulospinal axons as defined by EGFP-mRFP fluorescence colocalization (Fig. 1A-B). Tracking oligodendrocyte process-axon interactions by time-lapse imaging in control larvae revealed a wide range of cell-cell behaviors. These included formation of new reticulospinal axon interactions (gain), stable axon interactions, and retraction of axon interactions (loss) (Figure 1B).

**Figure 1.**
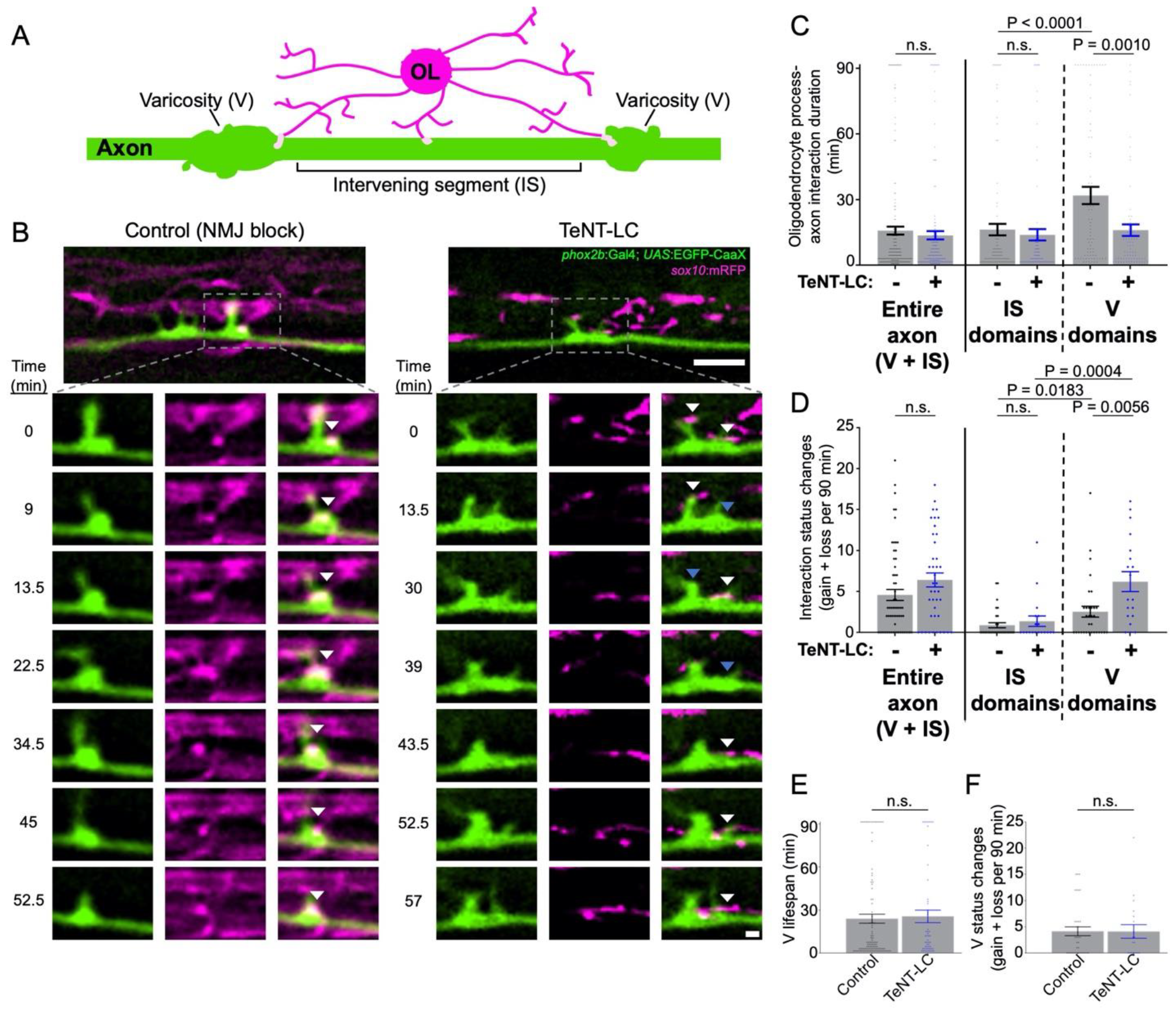
Synaptic vesicle release stabilizes oligodendrocyte interactions at axon varicosities. (A) Diagram illustrates an individual reticulospinal axon (green) and oligodendrocyte (OL) process extensions (magenta). Light pink shapes on the axon represent oligodendrocyte interactions. Axonal domains are segmented into varicosities (V) and intervening segments (IS). (B) Representative time-lapse images show oligodendrocyte-axon interactions over time in control (left) and TeNT-LC-expressing larval zebrafish (3 dpf). Lower inset images are individual timepoints derived from 90 minute time-lapse imaging sessions. Transgenic reporter expression of membrane-tethered fluorescent proteins label oligodendrocyte (magenta) and individual reticulospinal axon (green) membranes. White arrows indicate EGFP-mRFP colocalization (interaction sites) and blue arrows denote the location of previous colocalization (retracted interactions). Images are spinal cord lateral views with anterior left and dorsal up; scale bars are 5 μm (upper) and 1 μm (insets). (C) Scatter plot points represent oligodendrocyte-axon interactions and their duration in minutes. (D) Graph represents the total number of times oligodendrocyte-axon interactions were gained or lost at an individual reticulospinal axon segment across the entire 90 minute time-lapse. (E) Graph shows the length of time that individual axon varicosities persisted during the 90 minute time-lapse. (F) Scatter plot points show the number of times that varicosities and intervening segments changed identity at individual axon segments during time-lapse imaging. For C-F, bars represent mean ± SEM; sample size = 61 control and 38 TeNT-LC axon segments analyzed from 23 control and 14 TeNT-LC larvae. Reported P-values were obtained using the Mann Whitney test (C, D) or unpaired t-test (E, F).

To test the hypothesis that synaptic vesicle release regulates the temporal dynamics of oligodendrocyte-axon interactions, we used a loss of function approach to block synaptic vesicle release by mRNA-based tetanus neurotoxin light chain (TeNT-LC) overexpression. When initially tracking the duration, gain, and loss of oligodendrocyte-reticulospinal axon interactions over time, we observed no difference between control and TeNT-LC conditions (Figure 1C-D, entire axon). However, while acquiring and analyzing these data, we noted that interactions frequently formed at axon segments with increased local diameter (axon varicosities; V; Supplementary Fig. S1), and that interactions at varicosities appeared to have different temporal dynamics than those formed at neighboring, thinner axon segments (intervening segments; IS). We therefore separately assessed the behavior of interactions that occurred at varicosities and intervening segments. In control conditions, axon varicosities stabilized oligodendrocyte process interactions for a longer duration than interactions occurring at thin intervening axon segments (Fig. 1B-C). In contrast, TeNT-LC expression reduced the duration of oligodendrocyte-axon interactions specifically at varicosities (Fig. 1B-C). Comparatively, the durations of interactions at thin intervening axon segments in zebrafish larvae expressing TeNT-LC were indistinguishable from controls (Figure 1C).

We also tracked the frequency that axon segments underwent changes to interaction status, as defined by gain or loss of oligodendrocyte interactions. Blocking synaptic vesicle release increased the frequency that axon varicosities acquired and lost oligodendrocyte interactions (Figure 1B, D). In contrast, TeNT-LC expression did not alter oligodendrocyte-axon interaction dynamics at intervening axon segments (Fig. 1D). Taken together, these observations indicate that synaptic vesicle release is necessary to support prolonged interactions between oligodendrocyte processes and the varicosities of reticulospinal axons, but not at intervening axon segments.

One possibility is that synaptic vesicle release supports prolonged interactions by regulating the duration that axon varicosities remain in existence. Specifically, if varicosities disappeared more rapidly in TeNT-LC conditions, this could shorten oligodendrocyte-axon interaction durations by removing the favorable varicosity interaction sites. However, by measuring the duration that individual axon varicosities persisted, we detected no difference between control and TeNT-LC conditions (Fig. 1E). Additionally, the frequency that varicosities formed or converted into thinner intervening segments was also indistinguishable between groups (Fig. 1F). These data reject the notion that synaptic vesicle release dependent control of axon morphology itself is responsible for the observed changes to oligodendrocyteaxon interactions.

To further test the potential involvement of synaptic vesicles in oligodendrocyte-axon interactions, we next asked if synaptic vesicles are present at interaction sites. Expression of a Synaptophysin-EGFP (Syp-EGFP) fusion protein into TdTomato-CaaX expressing *phox2b*^*+*^ reticulospinal neurons showed EGFP^+^ puncta within and along the length of individual axons (Fig. 2A). Notably, we observed Syp-EGFP puncta in the majority of axon varicosities, but infrequently in intervening axon segments (Fig. 2B). To directly compare the location of Syp-EGFP puncta to oligodendrocyte processes, we next outcrossed *Tg(phox2b:Gal4), Tg(UAS:Syp-EGFP)* to *Tg(sox10:mRFP)*. We observed Syp-EGFP puncta in close proximity to oligodendrocyte processes at developmental timepoints corresponding to those used for time-lapse imaging in Figure 1 (Fig. 2C). To test the hypothesis that interactions occur preferentially at synaptic vesicle-enriched axon domains, we compared the density of *sox10*:mRFP^+^ process interactions at axon segments possessing or lacking Syp-EGFP fluorescent puncta. In support of this hypothesis, axon domains enriched with Syp-EGFP puncta showed an interaction density 2-fold greater than that observed at neighboring axon domains that lacked Syp-EGFP fluorescent puncta (Figure 2C-D).

**Figure 2.**
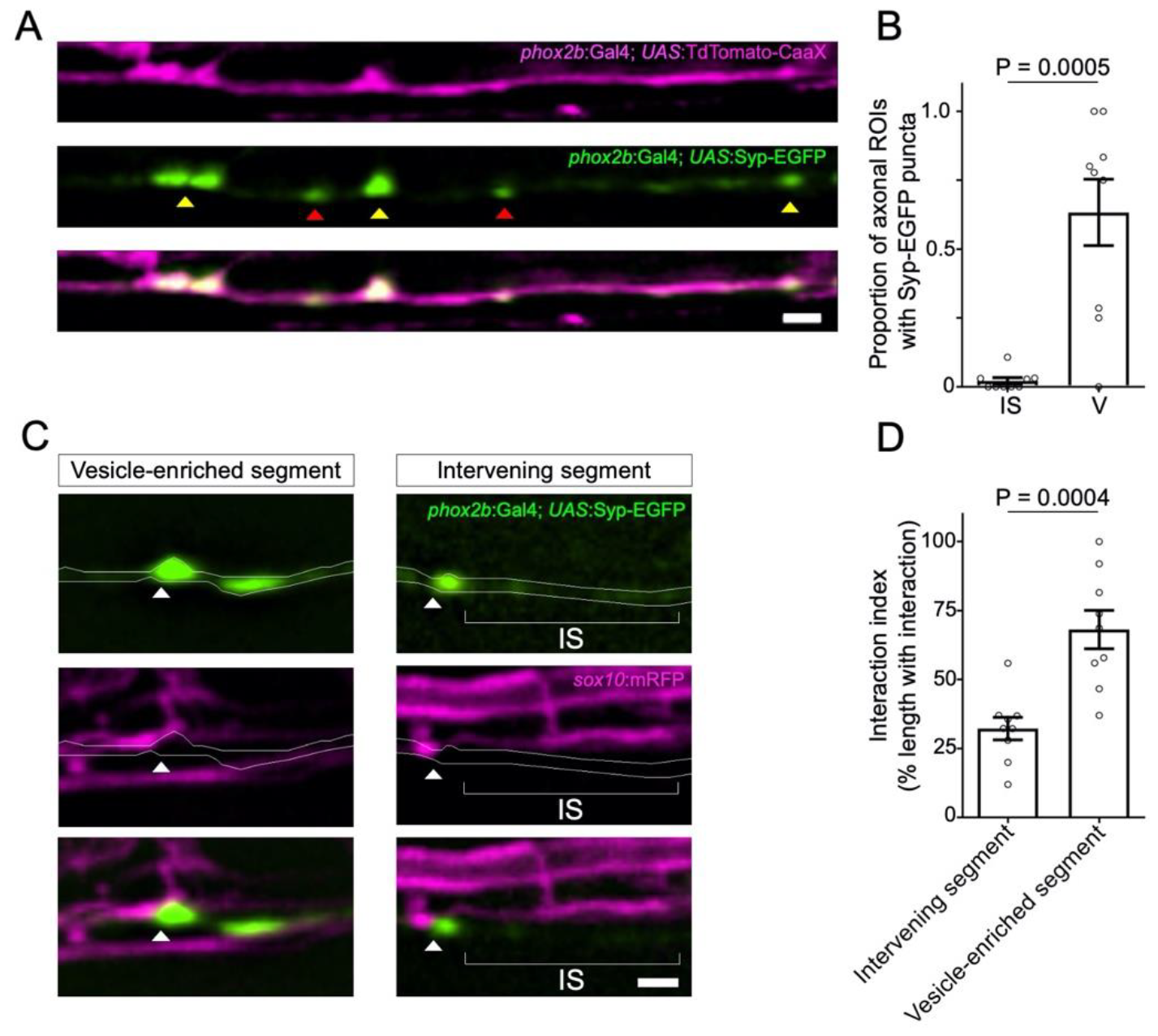
Syp-EGFP^+^ synaptic vesicles localize at sites of oligodendrocyte process interaction. (A) Representative confocal images of TdTomato-CaaX-labeled reticulospinal axons and EGFP-labeled synaptic vesicles in 4 dpf zebrafish spinal cord. Note Syp-EGFP puncta at both axon varicosities (yellow arrowheads) and intervening segments (red arrowheads). (B) Graph displays the proportion of axon ROIs that contained Syp-EGFP puncta. Each point represents the proportion of ROIs containing EGFP puncta per individual reticulospinal axon analyzed. Sample size = (number of ROIs, number of axons, number of animals) 339, 9, 8; bars show mean ± SEM, paired t-test. (C) Syp-EGFP and mRFP-labeled oligodendrocyte membranes observed in the larval zebrafish spinal cord (4 dpf). Arrowheads point to Syp-EGFP puncta associated with mRFP-labeled oligodendrocyte process extensions. (D) Graph displays the percent coverage of vesicle-enriched axon segments and intervening axon segments by oligodendrocyte process extensions. Sample size = (number of axons, number of animals) 9, 9; bars show mean ± SEM, unpaired t-test. For A and C, images are lateral view with anterior left and dorsal up; scale bars = 2 μm.

Does synaptic vesicle release stabilize interactions between oligodendrocyte processes and all axons, irrespective of neuronal subtype identity? If so, we predicted that blocking synaptic vesicle release would increase overall oligodendrocyte process motility while decreasing the frequency of process stalling behavior. To test this, we labeled individual oligodendrocytes with TdTomato-CaaX and tracked process motility by time-lapse imaging. Use of 15 second intervals between time-lapse frames captured rapid process movements and enabled tracking of individual process behaviors (Fig. 3A). In both control and TeNT-LC groups, we observed a similar frequency of process extension, retraction, and stall behaviors (Fig. 3A-B). In contrast, treatment with Latrunculin A, which blocks actin polymerization and disrupts cytoskeletal remodeling (positive control), increased the frequency of process stall behaviors, while decreasing the frequency of process extension or retraction behaviors (Fig. 3A-B). We also tracked individual processes frame-by-frame to evaluate the motility rate of process movements. Whereas the motility rate of process movements in control and TeNT-LC conditions were indistinguishable, Latrunculin A reduced process motility rate (Fig. 3A, C).

**Figure 3.**
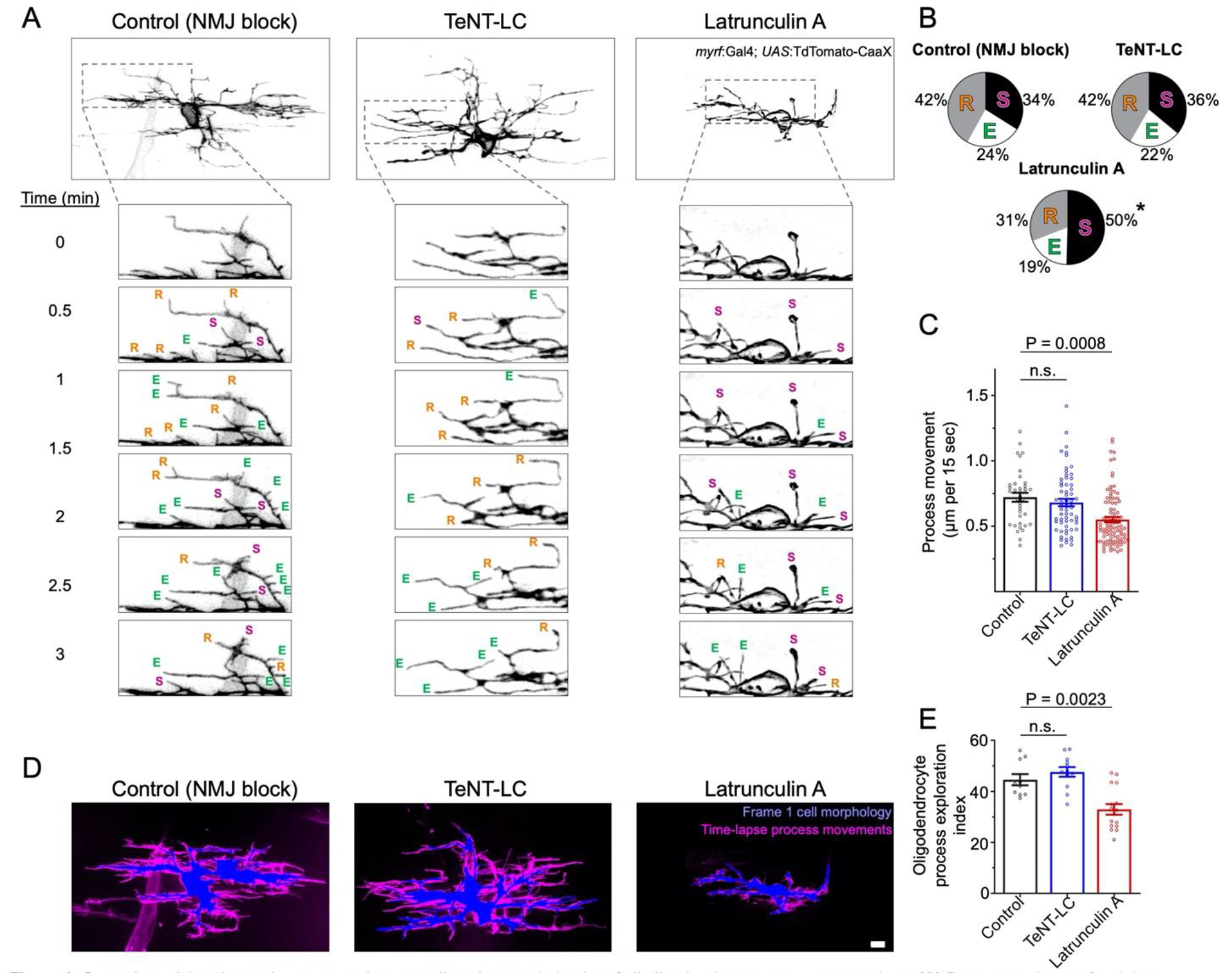
Synaptic vesicle release does not regulate overall exploratory behavior of all oligodendrocyte process extensions. (A) Representative confocal time-lapse images of TdTomato-CaaX labeled single oligodendrocytes (anterior left, dorsal side up). Within time-lapse series insets, E, R and S denote process extensions, retractions and stalls, respectively. (B) Pie charts display the proportion of processes that extended, retracted, or stalled during time-lapse imaging. Statistical testing (ordinary one-way ANOVA, Dunnett’s post-test) indicated increased stall behavior in the Latrunculin A (125 nM) group (P < 0.0001 for LatA vs control) but no change caused by TeNT-LC (P = 0.87 for TeNT-LC vs. control). (C) Graph depicts the frame-to-frame rate of process movements, with individual points representing the average distance that individual processes moved during each 15 second time-lapse interval (per cell). Sample size (tracked processes, cells) = 39, 8 (control), 60,8 (TeNT-LC), 106, 19 (Latrunculin A). (D) Composite images represent the extent of process movements during a 15 minute time-lapse imaging window. Blue shaded areas represent the morphology of individual oligodendrocytes at the time of the first time-lapse frame. Magenta shaded areas represent all pixels surrounding the original position where processes moved into or occupied for one or more frames during the time-lapse. This magenta shaded region is a maximum intensity over time projection, thereby representing the history of process movements surrounding the blue first frame position. Scale bar = 5 μm. (E) Summary of the percentage of the area (μm^2^) surrounding the blue first frame position that was occupied by processes during the time-lapse (magenta). Percentage of area measurements were restricted to a 1.5 μm band surrounding the blue first frame position. Each data point represents the average percentage of area within the 1.5 μm band occupied by processes of individual oligodendrocytes (per cell). Sample size (individual cells) = 10 (control), 12 (TeNT-LC), 15 (Latrunculin A). For C and E, bars represent mean ± SEM; Ordinary one-way ANOVA, Dunnett’s post test.

To further evaluate potential differences in overall process motility during time-lapse imaging, we analyzed time-lapse data sets to determine the extent that process extensions explored the area surrounding oligodendrocyte cell bodies. We first outlined the area of the cell body and process extensions at the onset of time-lapse imaging, and subsequently determined the proportion of the surrounding area that became occupied by motile processes during any time frame of the 15 minute timelapse. In control and TeNT-LC groups, motile processes explored a similar proportion of the surrounding area. In contrast, positive control larva treated with Latrunculin A showed a reduced extent of oligodendrocyte process exploratory behavior (Fig. 3D-E). Collectively, these findings reject the hypothesis that synaptic vesicle release blockade alters the motility or exploratory behavior of all oligodendrocyte process extensions.

We further reasoned that if synaptic vesicle release alters the behavior of all oligodendrocyte process extensions equally, then changes to oligodendrocyte process extension to axon interaction profiles would be equal across all axons regardless of neuronal subtype identity. To test this, we made use of reporter lines marking distinct neuron subpopulations with EGFP fluorophores. By crossing each neuronal reporter line to *Tg(sox10:mRFP)*, we then evaluated the extent that oligodendrocyte process-axon interactions covered the lengths of individual axons of distinct axon subtypes. Inhibiting synaptic vesicle release caused a net increase in the percentage length of individual *phox2b*^+^ reticulospinal axons covered by interacting oligodendrocyte processes (Fig. 4).

**Figure 4.**
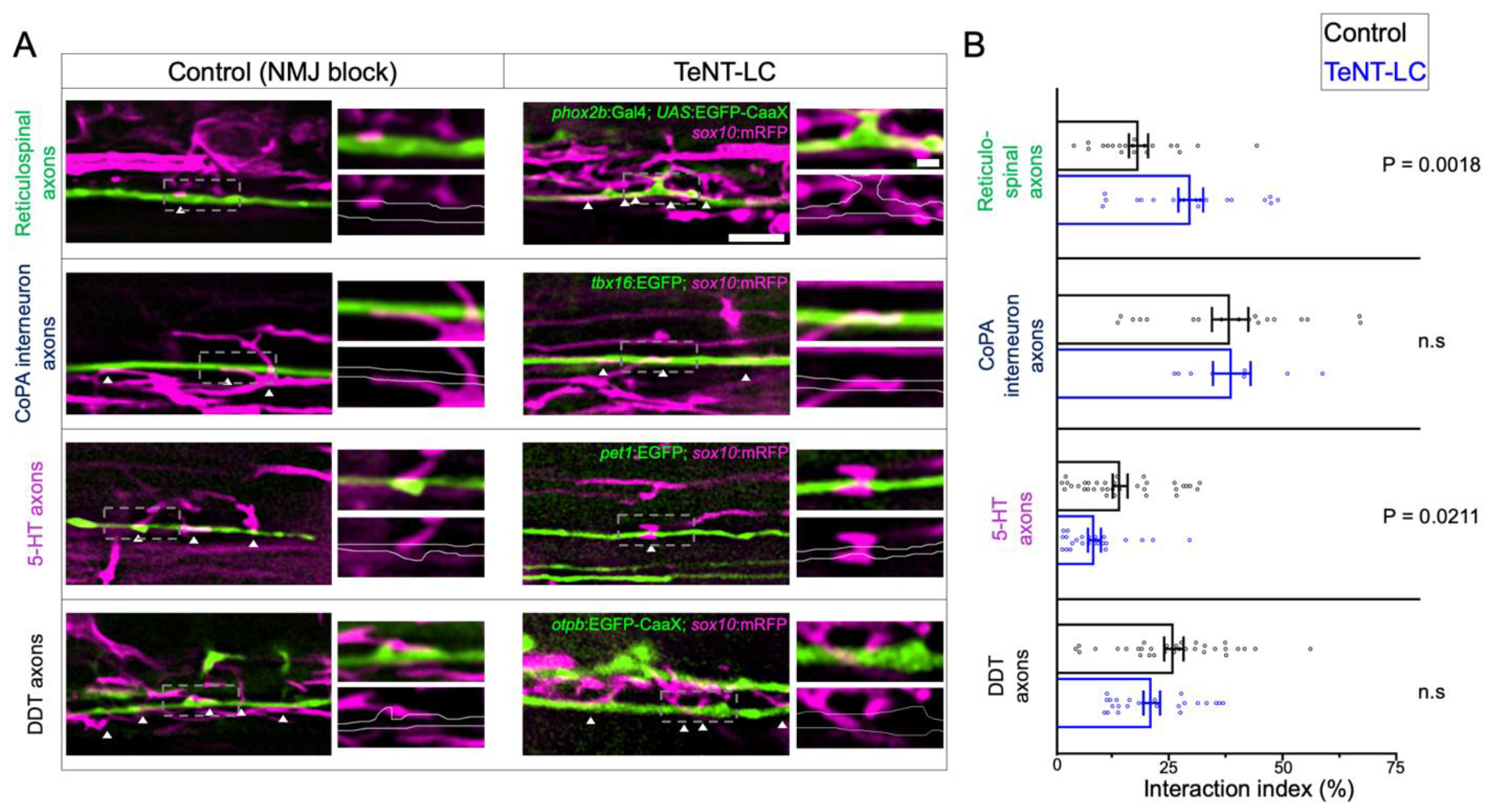
Synaptic vesicle release regulates oligodendrocyte-axon interactions in a neuron subtype specific manner. (A) Lateral view confocal images of GFP labeled axon interacting with oligodendrocyte processes labeled with RFP (anterior left, dorsal side up). White arrowheads point to sites of GFP-RFP colocalization. Scale bar = 5 μm, 1 μm (inset). (B) Graphs show the interaction index, which is defined as the percentage length of axon segments covered by interacting processes. Individual points represent the percent coverage of an individual axon. Sample size = number of animals, number of axons. Reticulospinal = 20, 20 (control), 19, 20 (TeNT-LC); CoPA = 18, 18 (control), 6, 8 (TeNT-LC); 5-HT = 19, 35 (control), 11, 24 (TeNT-LC); DDT = 23, 32 (control), 13, 24 (TeNT-LC). Black bars (controls) and blue bars (TeNT-LC) represent mean ± SEM. The P-values were derived using either a parametric t test (Reticulospinals, CoPA) or Mann-Whitney test (5-HT, DDT) based on whether data showed a normal distribution.

We next assessed interactions with axons of the dopaminergic diencephalospinal tract (DDT), which neighbor *phox2b*^*+*^ reticulospinal axons within in the ventral spinal cord medial longitudinal fasciculus (MLF). In contrast to our observations of *phox2b*^*+*^ reticulospinal axons, blocking synaptic vesicle release had no effect on the coverage of individual DDT axons with oligodendrocyte processes (Fig. 4). We also evaluated interactions between oligodendrocyte processes and two neuronal subtypes that extend axons in the dorsal longitudinal fasciculus. Similar to DDT axons, inhibiting synaptic vesicle release also had no effect on the interaction coverage of *tbx16*^+^ CoPA spinal interneuron axons. In contrast, blocking synaptic vesicle release decreased the extent that oligodendrocyte processes covered *pet1*^+^ serotonergic (5-HT) axons (Fig. 4). These findings indicate that synaptic vesicle release facilitates axon subtype-specific interactions with oligodendrocyte processes.

## Discussion

It is now well established that synaptic vesicle release facilitates axon-oligodendrocyte communication during myelination. Numerous studies together have established a model whereby vesicle release regulates myelin sheath growth and stabilization (Almeida et al., 2021; Hines et al., 2015; Hughes and Appel, 2019; Koudelka et al., 2016). Despite these advances, we still lack an understanding for how oligodendrocyte process extensions that have not formed myelin sheaths determine which axons to form interactions with, and how these interactions are stabilized. In this study we examined the role of synaptic vesicle release in oligodendrocyte process-axon interactions that precede initial axon wrapping and myelination. By labeling single oligodendrocytes with a membrane-tethered reporter, we found that the dynamic movements of the whole of an individual oligodendrocyte’s process arbor is not collectively sensitive to synaptic vesicle blockade. However, roles for synaptic vesicle release became apparent in our studies when we focused on interactions formed onto specific subsets of axons. Specifically, blocking synaptic vesicle release modified the density of processes interacting with reticulospinal and serotonergic axons, but not dopaminergic or commissural interneuron axons. In addition, focusing time-lapse measurements onto only the subset of process extensions interacting with reticulospinal axons revealed a role for synaptic vesicle release in stabilizing interactions. We therefore conclude that synaptic vesicle release regulates oligodendrocyte process extension dynamics and their interaction with individual axons in a neuron subtype-specific manner. Because we did not observe changes to process movements when examining all processes collectively, but did when examining those interacting with reticulospinal axons, we conclude that the behavior of oligodendrocyte process extensions can be modifiable by synaptic vesicle release when they interact with some but not all axons.

In our study, we found that axon varicosity domains play a unique role in the synaptic vesicle-mediated effects on oligodendrocyte-axon interactions. Varicosities, defined as segments of axons that appear swollen or of increased local diameter, were more likely to be interacted with by oligodendrocyte process extensions than were neighboring, thinner segments of axons (intervening segments). In alignment with other reports (Wake et al., 2015), we observed an accumulation of synaptic vesicles in axon varicosities. When we focused on sites of synaptic vesicle enrichment rather than axon morphology, these axon segments were similarly more likely to be interacted with by oligodendrocyte process extensions than were neighboring axon segments lacking synaptic vesicle enrichment. Consistent with a functional role for synaptic vesicle release at axon varicosities, we observed changes to interaction turnover and duration when synaptic vesicle release was blocked. Specifically, these effects were exclusive to interactions at axon varicosities and not observed at neighboring intervening axon segments, which were less likely to possess synaptic vesicle enrichment. However, a limitation to our study is that we did not specifically block synaptic vesicle release within neurons or oligodendrocyte-lineage cells. Therefore, while one possibility is that synaptic vesicle release from axons is responsible for stabilizing interactions, it is also possible that tetanus neurotoxin-sensitive vesicle dynamics within oligodendrocytes could be involved (Lam et al., 2022).

Though we focused on the pre-myelinating stage of oligodendrocyte development and process dynamics, our findings show some similarities to how synaptic vesicle release regulates myelinating oligodendrocyte processes. After nascent sheath formation, synaptic vesicle release prevents retractions and supports sheath growth (Almeida et al., 2021; Hines et al., 2015). However, these effects after initial myelination are not universal for all myelin sheaths, and instead depend on the specific axon and neuronal subtype being myelinated (Almeida et al., 2021; Koudelka et al., 2016). Similarly, when examining the effects of synaptic vesicle release on oligodendrocyte interactions with various defined neuronal subtypes, we observed that some but not all were regulated by synaptic vesicle release.

One question we did not address in our study is whether a subset of the interactions we labeled and tracked are neuron-OPC synapses. Since their discovery two decades ago, the temporal dynamics of neuron-OPC synapses have been unknown and only recently investigated (Bergles et al., 2000). By tracking the lifetime of PSD-95 and Gephryn in oligodendrocyte processes, Li et al. observed these complexes persisted on average 5-15 minutes, which is similar to the durations of interactions tracked in our study (Li et al., 2023). These findings by Li et al. demonstrate that oligodendrocyte process extensions containing postsynaptic proteins may form synapse-like interactions with axons that are transient in nature. An additional limitation of our study is that we did not address the long-term fates of oligodendrocyte-axon interactions. Instead, we designed our study to focus on the dynamics and spatial distribution of oligodendrocyte-axon interactions, and therefore used short time-lapse intervals (90 seconds) to more accurately track rapidly forming and disappearing interactions. A drawback inherent to time-lapse imaging is the limited number of images or timepoints that can be acquired before phototoxicity sets in, and to avoid this, we limited our time-lapse imaging sessions to 90 minutes. This trade-off did not enable us to follow interaction sites long enough to determine how frequently they developed into axon wrapping and myelination attempts. In contrast, Li et al. designed time-lapse experiments to track the fates of Gephryn-containing complexes in oligodendrocyte process for up to 16 hours. This long-term tracking revealed that Gephryn-containing complexes were predictive of where processes later matured into sheaths. Furthermore, the probability that Gephryn-containing processes formed sheaths was reduced by tetanus neurotoxin treatment, suggesting that synaptic vesicle release enables oligodendrocytes to determine which contacts should mature to form myelin sheaths (Li et al., 2023). In a separate study, Wake et al. reported that synaptic vesicle release blockade reduced local MBP translation and intracellular Ca2+ elevations that occurred within processes interacting at axon varicosities (Wake et al., 2015). Together with the results presented here, these studies collectively point to a model whereby synaptic vesicle release-mediated axo-glial communication may dictate where oligodendrocyte processes interact with axons and mature to form myelin sheaths.

## Materials and Methods

### Zebrafish lines and husbandry

All animal work performed in this study was approved by the Institutional Animal Care and Use Committee at Winona State University. Zebrafish embryos were raised at 28**°**C in egg water (0.0623 g/L Coralife marine salt) and staged according to hours post-fertilization or morphological criteria. Transgenic lines used in this study included *Tg(tbx16:EGFP)*^*uaa6/812c*^, *Tg(otpb*.*A:EGFP-CaaX)*^*zc49*^, *Tg(pet1:EGFP)*^*ne0214*^, *Tg(sox10:mRFP)*^*vu*^234, *Tg(nkx2*.*2a:EGFP-CaaX)*^*vu16*^, *Tg(phox2bb:EGFP)* ^*w37*^, *Tg(phox2bb:Gal4)*^*co21*^, *Tg(UAS:EGFP-CaaX)* ^*co18*^, *Tg(UAS:Sypb-GFP-2A-Tomato-CaaX)*^*rw0317*^, *Tg(UAS:Syp-EGFP)* (Hines et al., 2015), *Tg(UAS:TdTomato-CaaX)* (Faucherre and López-Schier, 2011), *Tg(myrf:Gal4)*^*win1*^ (Treichel and Hines, 2018).

### Genetic and pharmacological inhibitors

Tetanus neurotoxin light chain (TeNT-LC) mRNA microinjection was used to block synaptic vesicle release. pEXPR Tol2 SP6/CMV:TeNT-LC plasmid used as a template for mRNA synthesis was generated using Gateway LR cloning and plasmids from the Tol2 kit (Kwan et al., 2007). Entry and destination plasmids used included p5E SP6/CMV, pME-MCS, pME-TeNT-LC, p3E pA, and pDest Tol2 pA2. pME-TeNT-LC (no stop codon) middle entry was generated by sub-cloning the TeNT-LC coding sequence, lacking a stop codon, into pME-MCS plasmid. PCR amplification (primers 5’ GCTTGATTTAGGTGACACTATAGAATAC, 5’ ACTCCCGGGTTAAGCGGTACGGTTGTACAGG) was performed using Phusion High Fidelity DNA polymerase (New England Biolabs) using pCS2 TeNT-LC-GFP as a template (a gift from Martin Meyer). TeNT-LC coding sequence and pME-MCS backbone were digested using EcoR1 and Xma1 prior to ligation. Following gateway cloning to generate pEXPR Tol2 SP6/CMV:TeNT, mRNA synthesis (mMessage mMachine kit, Life Technologies), and phenol:chloroform purification, TeNT-LC mRNA (100-1000 ng/μl) was injected into 1-4 cell stage embryos. Prior to embedding for microscopy, a touch evoked response assay was conducted by tapping the head of larval zebrafish with a pin tool to ensure that each larvae imaged experienced total paralysis. No additional paralytic was used to perform microscopy in TeNT-LC-expressing embryos. Throughout the study, control larvae were paralyzed by neuromuscular junction (NMJ) blockade (pancuronium bromide, MP Biomedicals, 0.25 mg/mL or mivacurium chloride, LKT Laboratories, 0.5 mg/mL, in egg water). NMJ blockade was applied for 30 minutes prior to embedding in low melting-point agarose (1%, IBI Scientific), and embedded embryos were immersed in NMJ blockade solution during in vivo imaging. Treatment with Latrunculin A (Tocris) was performed in combination and concurrently with NMJ blockade (125 nM in egg water).

### In vivo imaging and analysis

Microscopy was performed using an Olympus IX81 equipped with a disk spinning unit (DSU), 60X 1.3 NA silicone oil immersion objective, LED illumination with narrow bandpass filters, and a Hamamatsu OrcaR2 CCD camera. Unless otherwise indicated, X-Y pixel sizes were 107.5 nm and confocal z-stacks were acquired at 400 nm internals. Larvae were embedded for lateral view and images acquired with dorsal up and anterior left. Image viewing, processing, background removal, and 3D deconvolution was performed within CellSens Dimension (Olympus, version 2.3).

Imaging of oligodendrocyte process extension interactions with axons was performed in 72-96 hpf larvae. For each transgenic reporter line, the same hemisegment range was selected and consistently viewed based on the following criteria: abundant RFP^+^ oligodendrocyte process extensions, incomplete myelination of Mauthner axons, and a maximum of 15 myelin sheaths within all z-stacks and the entire field of view. Oligodendrocyte process extension interactions with axons were identified by GFP-RFP colocalization within the same z plane. Percent interaction coverage was defined as the combined lengths of all interaction sites (on a singly-labeled axon) divided by the sum of the total length of the axon. In experiments where axons were marked by membrane-tethered fluorescent protein, axon varicosity domains were defined as local regions along the axon with diameter at least 15% greater than the immediately adjacent axon segments. Segmentation of axons into varicosity or intervening segment domains was performed by assigning consecutive 2 μm regions of interest, each spaced 2 μm apart, and identified as a varicosity or intervening segment using the criterion described above.

Imaging of Syp-GFP^+^ synaptic vesicles within reticulospinal axons was performed at identical developmental stages and at identical spinal cord hemisegments as oligodendrocyte-axon interaction experiments. Syp-GFP^+^ puncta were defined as regions of focal GFP fluorescence with 15% or greater fluorescence intensity than neighboring axon segments. The length of Syp-GFP^+^ and Syp-GFP^-^ axon segments were measured and the proportion (by length) that showed interactions with *sox10*:mRFP^+^ processes was determined by fluorescence colocalization.

Time-lapse tracking of oligodendrocyte process extension movements was performed by crossing *Tg(myrf:Gal4)* and *Tg(UAS:TdTomato-CaaX)* transgenes. Spinal cord hemisegments in 3 dpf larvae were selected and imaged when containing single, scatter-labeled oligodendrocytes with two or fewer nascent sheaths or sheath-like structures. Confocal z-stacks were acquired at 15 second intervals for 15 minute durations. Following image deconvolution in CellSens software, image stacks were imported into ImageJ. When needed, the Image Stabilizer and Bleach Correct plugins were used to correct for x-y drift and photobleaching between frames. A maximum intensity over z-projection was generated and process length and movements were manually tracked for each 15 second interval. To determine the extent that oligodendrocyte process extensions explored the surrounding area during the 15 minute time-lapse, the wand tool was used to first outline the morphology of the oligodendrocyte and its collection of process extensions in the first frame image. A band region of the 1.5 μm surrounding the initial morphology (from frame 1) was then generated as a new region of interest. Whereas no process extensions occupied this band region on frame 1 (by definition), motile processes generated fluorescence signal within this band region on subsequent time-lapse frames. The extent the band region was explored by process extensions during the time-lapse was determined by measuring the percentage area of the band region of interest with fluorescence signal on a maximum intensity over time projection image.

Oligodendrocyte process extension-axon interactions were tracked using time-lapse microscopy in 3 dpf larvae. Imaging was performed in spinal segments with singly-labeled *phox2b*^+^ axons, abundant *sox10*:mRFP^+^ oligodendrocyte process extensions, incomplete myelination of Mauthner axons, and no more than 15 myelin sheaths adjacent to the axon and within the z-planes acquired. Images were acquired at 1.5 minute intervals for 90 minutes with 0.4 μm z-intervals and 0.215 μM pixel size. Following image deconvolution, X or Y drift was corrected using the Image Stabilizer plugin within ImageJ. To track interactions, axons were divided into separable regions of interest (2 x 2 μm) for time-lapse analysis that were sub-categorized as intervening axon segments or as containing an axon varicosity. Fluorescence colocalization within the same z-plane and timepoint was used as an indicator of interaction. Gain or loss of interaction was defined as the appearance or disappearance of fluorescence colocalization with respect to the previous frame (time). Interaction duration was determined by summing the number of consecutive time-points an ROI contained an interaction.

### Quantification and Statistical analysis

Statistical analyses and graphs were generated using Prism 7.05 (GraphPad) or Microsoft Excel. Paired and unpaired student’s t-tests were used for datasets with normally distributed data between control and treatment groups. For comparisons between samples with non-normal distributions, Mann-Whitney tests were conducted. For comparisons between 3 or more groups, ordinary one-way ANOVA tests were used. Variance within datasets were represented as standard error of means.

### Availability of data and materials

All datasets, zebrafish lines, plasmids, and other reagents are available upon reasonable request.

## Acknowledgements

This work was supported by National Science Foundation CAREER award IOS-1845603 (JHH), National Multiple Sclerosis Society RG-5274A1/T and PP-1706-28071 (JHH). National Multiple Winona State University Special Project awards (251.0225, 251.0253, 251.0300, 251.0327), Minnesota State Colleges and Universities Professional Improvement Funds and Leveraged Equipment Funds, and Winona State University Undergraduate Research & Creative Projects grants (MM, TH, AT). We thank Erika Vail, Mary Diekmann, Amy Runck, Richard Deyo, Scott Segal, Scott Steele, Silas Bergen, Bailey Otto, Madison Schaefer, Shelbey Strandberg, John Butrum, Josh Giese, Anthony Gates, Katie Waller, and Isaiah Murray for technical assistance and valuable feedback. We also thank Mark Masino, Bruce Appel, Hernan Lopez-Schier, the Zebrafish International Resource Center (ZIRC), and the National BioResource Project (NBRP) for providing zebrafish lines.

