## Supplementary Figure S1 for "Synaptic vesicle release regulates pre-myelinating oligodendrocyte-axon interactions in a neuron subtype-specific manner"

***Supplementary Figures***

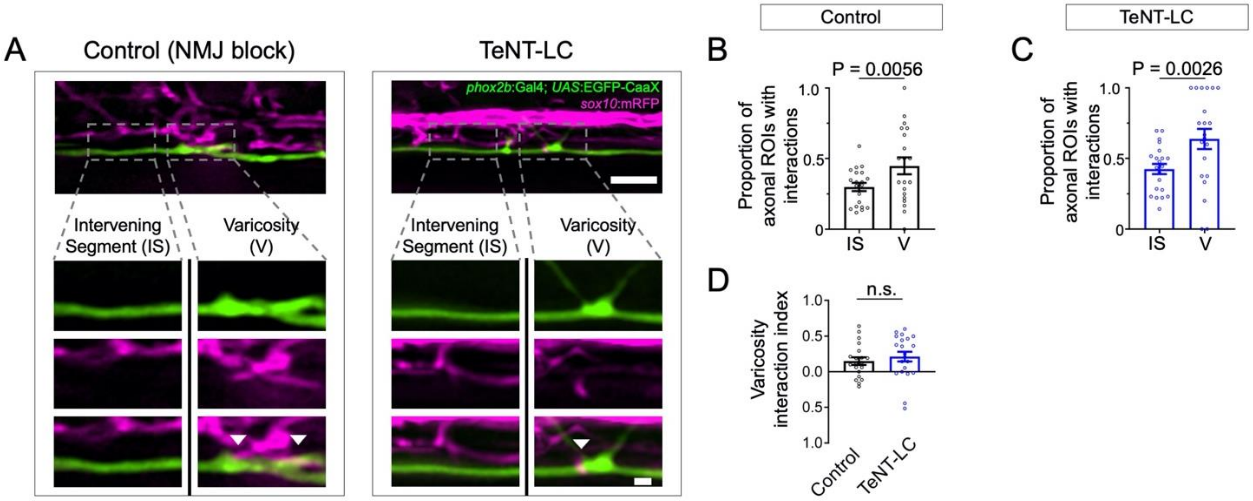


**Supplementary Figure S1. Oligodendrocyte processes display spatial preference for interactions with axon varicosities.** (A) Lateral view representative confocal images showing EGFP-CaaX-labeled reticulospinal axons and mRFP-labeled oligodendrocyte processes in control and TeNT-LC expressing larvae (4 dpf). Arrowheads point to colocalization. Images show lateral view, anterior left, dorsal up; scale bars represent 5 µm (upper) and 1 µm (inset images). (B-C) Graphs display the proportion of 2 µm ROIs that contained an oligodendrocyte process extension-axon interaction for either intervening axon segments or varicosities. Individual scatter points represent the proportion of ROIs with an oligodendrocyte process extension-axon interaction for a single axon. (D) Graph summarizes the extent that interactions are enriched at varicosities. Varicosity interaction index is the proportion of varicosities with interactions minus the proportion of intervening segments with interactions for each individual axon, which are shown as individual points on the scatter plot. For B-D, bars represent mean ± SEM; Control sample size = (number of ROIs, number of axons, number of animals) 569, 20, 20. TeNT-LC sample size = 537, 21, 20. Reported P-values were obtained using paired t-test (B, C) or unpaired t-test (D).
